# Phosphoprotein-based Biomarkers as Predictors for Cancer Therapy

**DOI:** 10.1101/675637

**Authors:** Angela M. Carter, Chunfeng Tan, Karine Pozo, Rahul Telange, Roberto Molinaro, Ailan Guo, Enrica De Rosa, Jonathan O. Martinez, Shanrong Zhang, Nilesh Kumar, Masaya Takahashi, Thorsten Wiederhold, Hans K. Ghayee, Sarah C. Oltmann, Karel Pacak, Eugene A. Woltering, Kimmo J. Hatanpaa, Fiemu E. Nwariaku, Elizabeth G. Grubbs, Anthony J Gill, Bruce Robinson, Frank Gillardon, Sushanth Reddy, Renata Jaskula-Sztul, James A. Mobley, M. Shahid Mukhtar, Ennio Tasciotti, Herbert Chen, James A. Bibb

## Abstract

Disparities in cancer patient responses have prompted widespread searches to identify differences in sensitive vs. non-sensitive populations and form the basis of personalized medicine. This customized approach is dependent upon the development of pathway-specific therapeutics in conjunction with biomarkers that predict patient responses. Here, we show that Cdk5 drives growth in subgroups of human patients with multiple types of neuroendocrine neoplasms. Phosphoproteomics and high throughput screening identified novel phosphorylation sites downstream of Cdk5. These phosphorylation events serve as biomarkers of Cdk5 activity and effectively pinpoint Cdk5-driven tumors. Toward achieving targeted therapy, we demonstrate that mouse models of neuroendocrine cancer are responsive to selective Cdk5 inhibitors and biomimetic nanoparticles are effective vehicles for enhanced tumor targeting and reduction of drug toxicity. Finally, we show that biomarkers of Cdk5-dependent tumors effectively predict response to anti-Cdk5 therapy in patient-derived xenografts. Thus, a phosphoprotein-based diagnostic assay combined with Cdk5-targeted therapy is a rational treatment approach for neuroendocrine malignancies.

---

Cyclin dependent kinases (Cdks) are a family of proline-directed serine/threonine kinases that are required for progression of normal cell division. Typical Cdks are regulated through binding to cyclins, proteins which are expressed at varying levels at distinct stages of the cell cycle^1^. As master regulators of cell division, Cdk1/2/4/6 currently serve as popular targets for cancer therapy development^2–4^. Although Cdk5 shares ~60% sequence identity with founding family members Cdk1 and Cdk2^5^, it possesses non-canonical features. Specifically, Cdk5 is not required for normal cell cycle progression and is not activated by cyclins. Cdk5 is regulated instead through interactions with cofactors p35 and p39^6^. Although Cdk5 is expressed in a broad number of tissues^5^, its activators are mostly restricted to neuronal cells^7^ where Cdk5 activity is important for CNS development and cognitive processes such as learning and memory^8^. Cleavage of p35 and p39 by calpain produces truncated activators, p25 and p29, respectively^9^. These cleavage products have increased protein stability and mislocalize in cells due to removal of an N-terminal myristoylation site^10^. In neurons, p25 aberrantly activates Cdk5 and has been linked to neurotoxicity, neuronal injury, and neurodegeneration^6^.

Recent research suggests the aberrant activation of Cdk5 in non-neuronal cells can usurp signaling components involved in the cell cycle to drive proliferation^11^. Expression of Cdk5 and p35/p25 has been shown for three types of neoplasms originating from neuroendocrine (NE) cells: medullary thyroid carcinoma (MTC)^12^, small cell lung cancer (SCLC)^13^, and pituitary adenomas^14^. In MTC, inhibition of Cdk5 activity decreases rates of cell growth^12,15^; in SCLC and pituitary adenomas, it decreases migration and invasion^16,17^. Expression of p25 in thyroid C-cells produces MTC in mice^12^, in part through alteration of traditional cell cycle regulatory components^18^. Here we show that Cdk5 and p35/p25 expression may be an important driver of many types of NE cancer and that aberrant Cdk5 activity allows for a diagnostic-coupled treatment strategy that targets this protein kinase.

## Cdk5 in NE Neoplasms

To better understand the potential role of Cdk5 across multiple forms of NE malignancies, we assessed expression of the kinase and its activating cofactors in different NE tumor and cancer types. Histological analysis demonstrated that Cdk5 and p35/p25 are present throughout various human NE neoplasms including MTC, pheochromocytoma (PHEO), pituitary adenoma, SCLC, pancreatic NE tumors, and gastrointestinal NE tumors (Fig. 1a). Furthermore, these proteins are present in cell lines derived from multiple types of NE neoplasms, including three human MTCs (TT, MTC-SK, and SIN-J), a human progenitor PHEO (hPheo1)^19^, a human pancreatic carcinoid (BON), a rat insulinoma (INS), and two human SCLCs (H146 and H1184, Fig. 1b). Selective Cdk5 inhibition by Indolinone A (Indo A) blocked all human NE cancer cell growth more potently than it affected normal human fibroblasts or rat INS cells (Fig. 1c and Ext. Data Fig. 1a-h). Interestingly, the aberrant Cdk5 activator, p25, was present in all human cell lines derived from naturally occurring tumors but not the rat line generated using irradiation^20^ (Fig. 1b). Cells treated with Cdk5 inhibitors typically showed flattening and smoothing of cell body consistent with a less malignant phenotype (Fig. 1d).

**Fig. 1:**
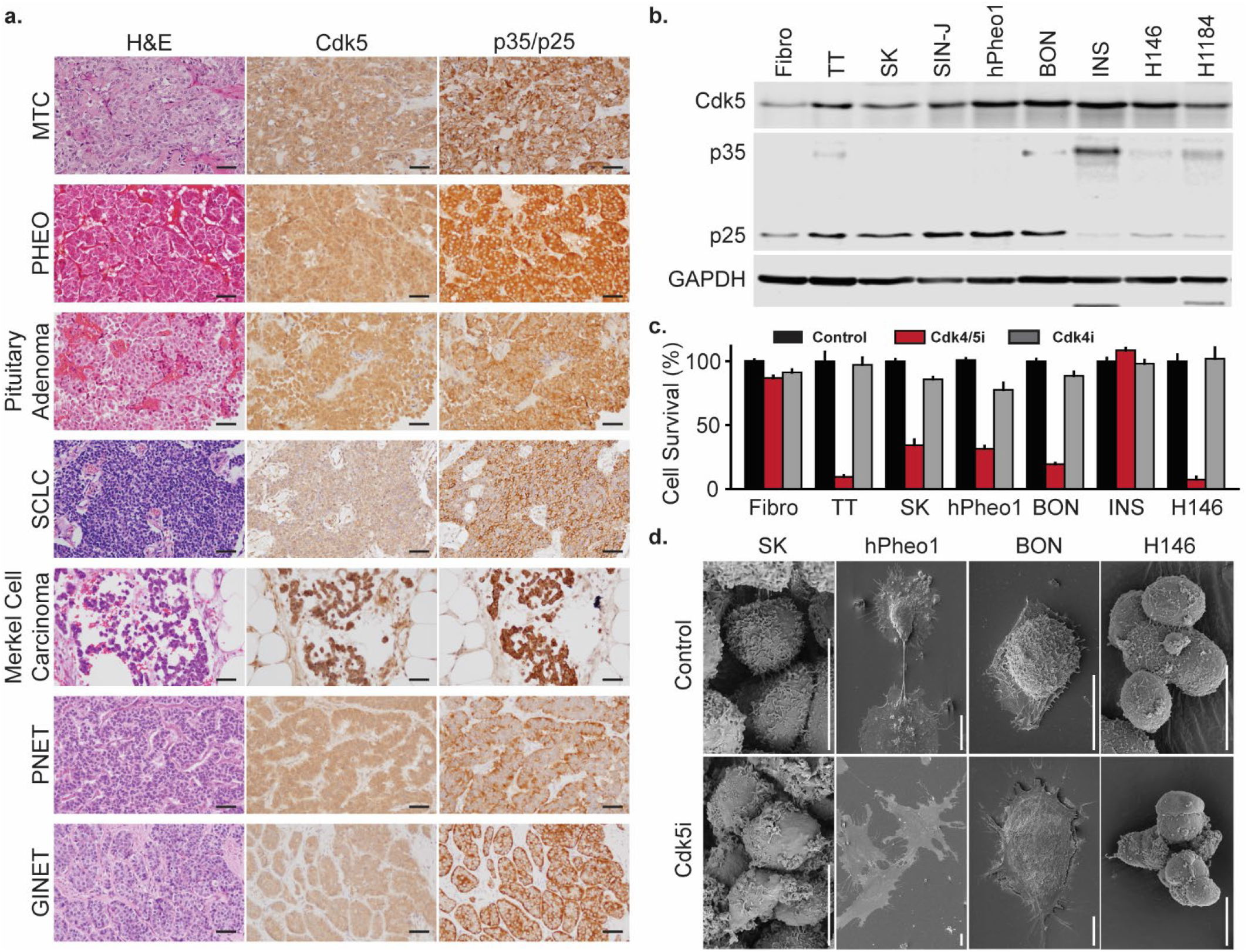
Cdk5 promotes growth in human NE tumors. **a**, H&E stain, IHC for Cdk5, and IHC p35/p25 in human NE tumors. Scale bars = 50 μm. **b**, Immunoblot of Cdk5 pathway components in fibroblasts and NE cells. **c**, Fibroblasts and NE cell lines were treated with 0.02 % DMSO (Control), 0.3 μM Indo A (Cdk4/5i), or 0.3 μM Indo B (Cdk4i) and monitored for effects on cell growth. Error bars represent SEM. (Full curves and IC_50_ values are reported in Ext. Data Fig. 1). **d**, Scanning electron microscopy of NE cells treated with Control (0.02% DMSO) or Cdk5i (hPheo1 and BON – 2 μM Indo A; MTC-SK and H146 – 5 μM CP6813101) for 4 h. Scale bars = 10 μm.

Indo A inhibits Cdk5 activity with high affinity, but also affects Cdk4^21^. Importantly, the structurally related Cdk4-specific inhibitor, Indo B, was 6.4-15.4 fold less potent at preventing human NE cell proliferation (Fig. 1c and Ext Data Fig. 1a-h). Growth of multiple NE cancer cell lines was likewise blocked by the broad spectrum Cdk inhibitors Roscovitine and Dinaciclib. CP681301, a selective inhibitor of both Cdk2 and Cdk5, also blocked growth of NE cancer cells whereas the Cdk2 specific inhibitor CVT313 had a greatly reduced effect (Ext. Data Fig. 1i-l). Thus, Cdk5 inhibition was necessary to robustly inhibit proliferation of cells derived from multiple forms of NE neoplasms. These data indicate that Cdk5 and p35/25 expression characterize at least a portion of all NE neoplasms and aberrant Cdk5 activity is a major contributor to the growth of NE cancer cells.

## Downstream targets of Cdk5

To determine which pathways Cdk5 may target to drive NE cell proliferation, we used a unique bi-transgenic mouse model of MTC (NSE-p25OE mice), developed in our laboratory, in which tumors arise at the natural organ site in the presence of an intact immune system. These mice express an activator of Cdk5, p25-GFP, in C-cells of the thyroid under the control of a doxycycline (dox) regulatable promotor (Fig. 2a)^12^. This system can be used to generate actively growing MTC tumors through expression of p25-GFP as well as growth-arrested MTC tumors through initial expression and subsequent suppression of p25-GFP via readministration of dox (Fig. 2b-c). As C-cells only comprise 3% of a normal thyroid, this system allows generation of sufficient quantities of C-cells for direct comparison of the signaling state between dividing and non-dividing populations.

**Fig. 2:**
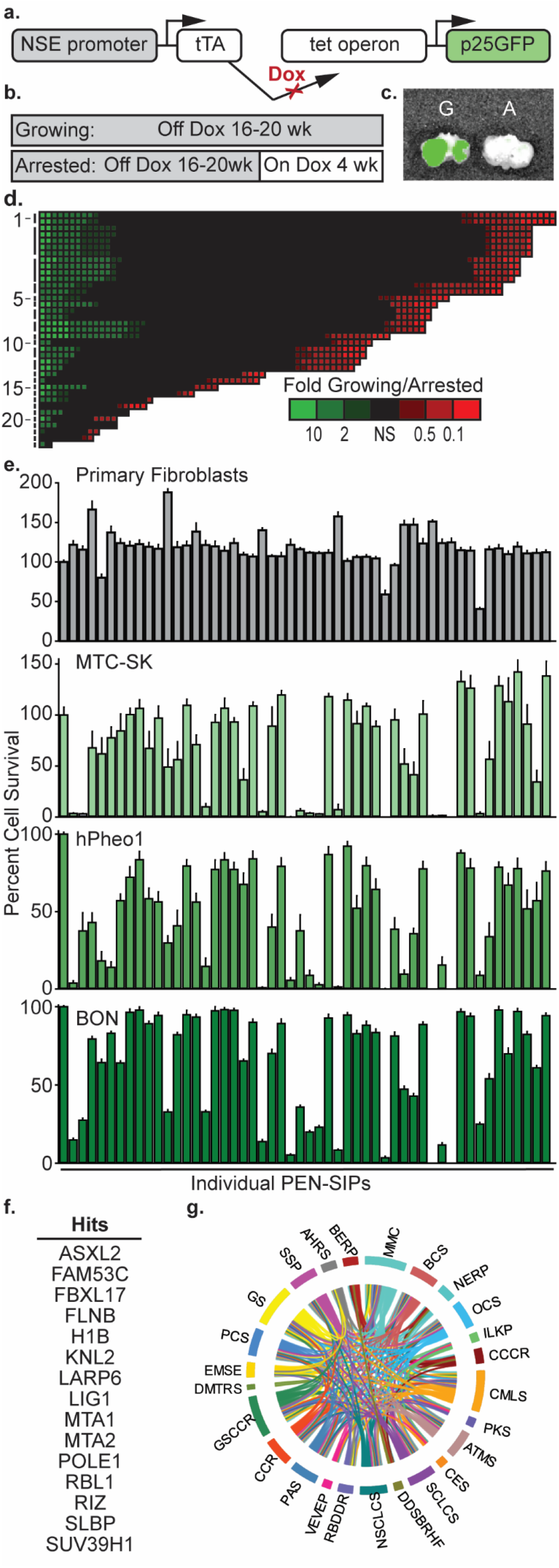
Identification of potential tumorigenic phosphoprotein signaling pathways. **a**, Schematic of bi-transgenic system for regulated tissue specific expression of p25GFP. **b**, Diagram of induction paradigm for generation of growing and arrested tumor tissue. **c**, Overlay of photograph and GFP fluorescence from IVIS analysis of resected trachea/esophagus with bilateral growing (G) and arrested (A) tumor tissue. **d**, Phosphoproteome of mouse MTC tumors represented as a heat map of phosphopeptide levels in growing versus arrested tumors sorted by protein group as indicated in Ext. Data Table 1 (y – axis) and fold change intensity (x – axis). **e**, Growth/survival assay of NE cell lines treated with control (0.3 % DMSO) or the PEN-SIPs (30 μM) indicated in Ext. Data Table 2. Error bars represent SEM. (n=7-8). **f**, Hits selected from (e). **g**, Ingenuity Pathway Analysis of hits from **e**.

Phosphoproteomic analysis of growing versus arrested MTC tumors, using PhosphoScan technology^22^, revealed global differences in the proline directed S/T phospho-signaling network including over 250 peptides with elevated phosphorylation levels in growing tumors (Fig. 2d and Ext. Data Table 1). From this set of phosphorylation sites, those not conserved in humans or conforming to a stringent Cdk5 phosphorylation sequence (S/T-P-x-K/H/R) were eliminated. From the remaining proteins, 50 of the most highly upregulated phosphorylation sites, with preference for those with established or suggested roles in cancer, were selected for investigation as potential tumorigenic regulators (Ext. Data Table 2). Short interfering peptides (SIPs) containing the phosphorylation site flanked by 8 amino acids on both the N- and C-terminal sides were designed to selectively interfere with phosphorylation or function of the 50 targets. A cell penetrating sequence (RQIKIWFQNRRMKWKK) from penetratin (PEN) was added to the N-terminus of peptides to facilitate entry into cells. A high-throughput proliferation-based assay identified 15 SIPs that inhibited growth of NE cancer cells but not normal primary fibroblasts (Fig. 2e-f and Ext. Data Table 2), indicating that these novel sites were important for NE cancer cell growth.

We performed Ingenuity Pathway Analysis (IPA) to ascertain major signaling cascades and pathways that are associated with these 15 proteins. Among the predicted 25 statistically enriched canonical pathways, cell cycle regulation, DNA repair, and diverse cancer signaling pathways are a predominant feature (Fig. 2f and Ext. Data Table 3). Thus, the downstream targets identified here are associated with common cancer mechanisms.

## Biomarkers of Cdk5 activity

Phosphorylation state-specific antibodies were generated for detection of 6 novel sites (Ser608 ASXL2, Thr143 FAM53C, Thr709 FLNB, Thr202 LARP6, Ser110 KNL2, and Ser988 RBL1) and two sites previously identified as targets of Cdk2 (Ser17 H1B^23,24^ and Ser391 SUV39H1^25^) (Ext. Data Fig. 2). Phosphorylation of these sites, as well as the thoroughly established Ser807/Ser811 sites on RB^26,27^, was confirmed in mouse MTC tumors, in which growth was driven by expression of p25-GFP, and reduced in arrested tumors (Fig. 3a). In agreement, phosphorylation levels of 6 of these sites were dose-dependently decreased by the Cdk4/5 inhibitor Indo A in hPheo1 cells (Fig. 3b). Similar decreases in these phosphorylation sites were observed in human MTC-SK, TT, and BON cells (Ext. Data Fig. 3). This effect appeared Cdk5-specific as addition of the Cdk4 inhibitor Indo B to multiple NE cell lines had significantly less effect on the phosphorylation states of Thr143 FAM53C, Thr709 FLNB, Ser17 H1B, and Thr202 LARP6 compared to Indo A. In contrast, phosphorylation of Ser807/Ser811 RB, a known target of multiple Cdks, was decreased upon treatment with both inhibitors. Interestingly, the phosphorylation of the RB family member, RBL1, was more responsive to Indo A than Indo B in TT cells whereas phosphorylation of Ser391 SUV39H1 was more responsive to Indo A than Indo B in BON cells (Ext. Data Fig. 3). Overall, these data demonstrate that phosphorylation of these 6 proteins is dependent upon Cdk5 activity and suggests that these phosphorylation sites could serve as biomarkers of many types of Cdk5-driven NE tumors.

**Fig. 3:**
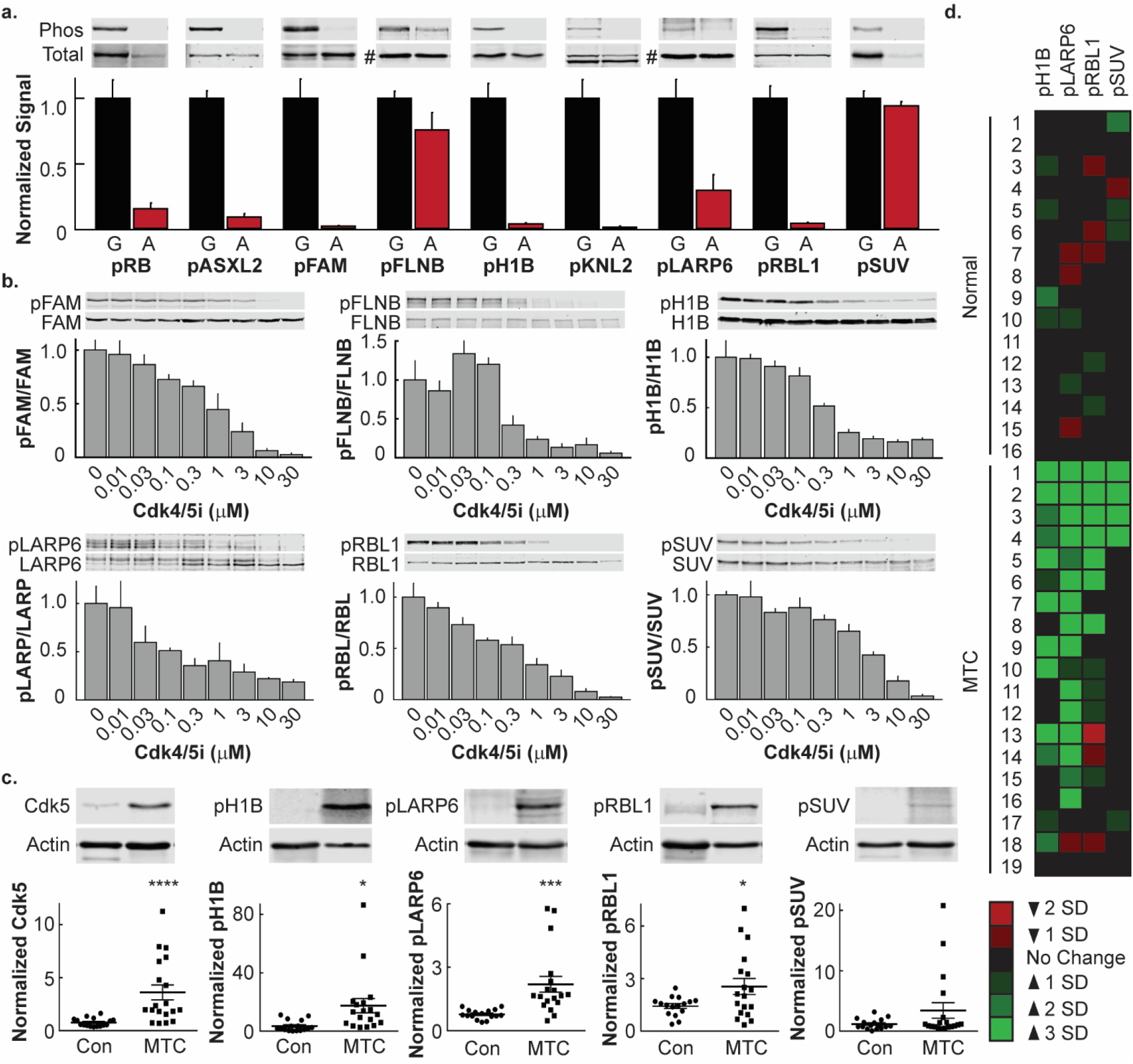
Phosphoproteins are biomarkers of Cdk5 dependent NE tumors. **a**, Immunoblot analysis of phosphoproteins in growing (G) and arrested (A) mouse MTC tumors. For pFLNB and pLARP6, total blot is actin (#); all others are normalized to each specific protein. (n=3-4). **b**, Immunoblot analysis of phosphoproteins in human hPheo1 cells treated with increasing concentrations of Indo A (Cdk4/5i) for 4 hours. (n=3-4 for all except 0.01, 1, and 30 μM points of pLARP6 where n=2) c, Immunoblot analysis of Cdk5 and phosphoproteins in normal human thyroid tissue and human MTC tumors. Due to sample size, samples were processed on three gels, each containing a reference for normalization. **d**, Heat map representation of immunoblot analysis of phosphoproteins from **c**, relative to the average and standard deviation (SD) of the total normal thyroid population, grouped by individual patient sample. All error bars represent SEM.

To determine if these Cdk5-dependent biomarkers occurred in human NE tumors, cohorts of MTC patient tumors and normal human thyroid tissues were compared. Immunoblot analysis revealed that three of the phosphorylation sites, Ser17 H1B, Thr202 LARP6, and Ser988 RBL1, were significantly elevated in the total tumor population and exhibited a positive correlation with overall Cdk5 expression levels (Fig. 3c and Ext. Data Fig. 4). The phosphorylation state of Ser391 SUV39H1 was only increased in a small portion of tumor samples but retained a positive correlation with Cdk5 expression. Distribution of all four phospho-sites varied across patients with phospho-LARP6 being the most commonly detected followed by phosphorylation of H1B and RBL1, respectively (Fig. 3d). Although 73% of patient tumors exhibited elevated Cdk5 levels, only 21% presented with elevation of all four biomarkers of Cdk5 pathway activity, emphasizing the fact that presence of a protein does not correlate 100% with function of that protein and highlighting the need for direct readouts of pathway activity such as phosphorylation of downstream substrates. Thus, Cdk5-dependent tumorigenic signaling may be considered patient-specific. Furthermore, detection of all four biomarkers in a patient could predict positive response to a Cdk5-targeted therapeutic approach.

## Cdk5 inhibitors as effective therapeutics in NE models

To evaluate the efficacy of anti-Cdk5 therapy when administered systemically, multiple animal models of NE cancer were used. First, the transgenic model of MTC driven directly by activation of Cdk5, NSE-p25OE, (Fig. 4a-b) was treated with vehicle or Indo A (10-30 mg/kg body weight, BW) once every three days for two weeks. Tumor growth was significantly reduced by 20 and 30 mg/kg of Indo A compared to control animals. Plasma levels of CEA, a marker of MTC^28^, were also reduced in treated animals, indicating therapeutic efficacy (Fig. 4c). As a second model, TT cells derived from a familial case of human MTC were used to generate xenograft tumors in nude mice. As with the transgenic model, Indo A attenuated TT cell tumor progression (Fig. 4d-e). While CEA was not detected in the plasma of these animals, levels of chromogranin A (ChA), another marker of MTC^29^, were reduced (Fig. 4f). Thus, Cdk5 inhibition blocks the growth of tumors in distinct models of MTC. The efficacy of Indo A was also assessed in a xenograft tumor model generated from human pancreatic NE tumor derived BON cells (Ext. Data Fig. 5). Indo A was equally effective in impeding the growth of these tumors, suggesting that targeted Cdk5 therapy could be effective across multiple NE cancer types.

**Fig. 4:**
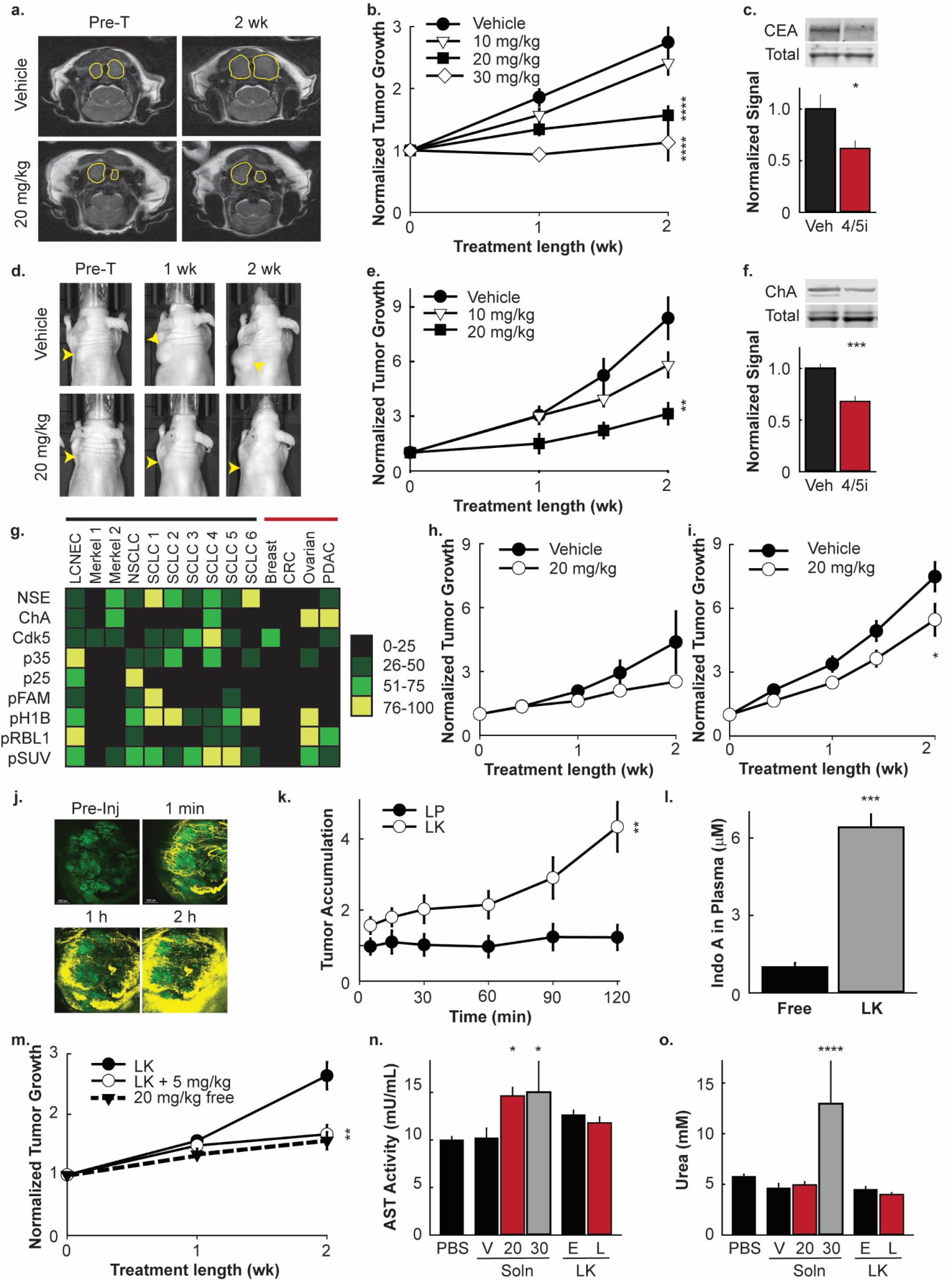
Targeted Therapy. **a**, Representative MRIs of NSE-p25 MTC model mice prior to treatment (Pre-T) or treated for 2 wk with vehicle or 20 mg/kg BW Indo A. Tumors outlined in yellow. **b**, Quantitation of tumor growth over time from MRI of NSE-p25 MTC mice treated with vehicle (n=6) or 10 mg/kg (n=4), 20 mg/kg (n=6), or 30 mg/kg (n=3-4) BW Indo A. **c**, Immunoblot analysis of blood plasma in vehicle and 20-30 mg/kg BW Indo A animals from **b. d**, Representative images of TT cell xenograft MTC model mice treated for 2 wk with vehicle or 20 mg/kg BW Indo A. Tumors marked by yellow arrowheads. **e**, Quantitation of tumor growth over time from caliper measurements of TT cell xenograft MTC model mice treated with vehicle or 10 mg/kg or 20 mg/kg BW Indo A (n=5-7). **f**, Immunoblot analysis of blood plasma in vehicle and 20 mg/kg BW Indo A animals from **e. g**, Heat map of phosphoprotein population percentile ranking from immunoblot analysis of NE tumors (black bar) and non-NE tumors (red bar) from PDX model mice. **h-i**, Quantitation of tumor growth over time from biomarker negative (**h**; n=5-6) and biomarker positive (**i**; n=8) PDX model mice treated with vehicle or 20 mg/kg BW Indo A (20 mg/kg). **j**, Representative images of MTC tumors from NSE-p25 mice injected with Cy5.5 labeled LKs and imaged *in vivo* over 2 h using IVM; green – tumor cells, yellow – LKs. **k**, Quantitation of accumulation of Cy5.5 labeled LPs and LKs in MTC tumors of NSE-p25 mice over time normalized by total tumor size (n=7-8). **l**, HPLC-MS analysis of Indo A in blood plasma from NSE-p25 MTC mice treated with 5 mg/kg BW Indo A (Free) or 5 mg/kg BW Indo A encapsulated in LKs (LK) (n=3) **m**, Quantitation of tumor growth over time from MRI of NSE-p25 MTC mice treated with empty LKs or 5 mg/kg BW Indo A encapsulated in LKs (LK + 5 mg/kg) compared to 20 mg/kg BW free Indo A from **b** (n=5-6). **n-o**, Analysis of blood plasma for AST activity (**m**) and urea level (**n**) in animals treated with PBS, Indo A in solution (V-vehicle alone, 20 mg/kg BW, or 30 mg/kg BW), or Indo A encapsulated in LKs (E-empty LK, L-drug loaded LK). All error bars represent SEM.

## Biomarkers are predictive of response to anti-Cdk5 therapy

We hypothesized that the phosphorylation states of a newly identified set of proteins could serve as biomarkers of Cdk5-driven tumors. If true, biomarker positive tumors should be responsive to Cdk5 inhibitor therapy while biomarker negative tumors should be non-responsive. To test this, a cohort of tumors from patient-derived xenograft (PDX) models of mixed origins was analyzed quantitatively for the presence of NE markers, Cdk5 pathway components, and putative Cdk5-dependent biomarkers (Fig. 4g). A Large Cell NE Carcinoma (LCNEC) model was identified that exhibited NE markers, expressed Cdk5 and high levels of its activators, and the 4 biomarkers. In contrast, a Merkel cell model (Merkel 2) was identified that expressed NE markers, but low Cdk5 pathway components, and low biomarker levels. Cohorts of these two disparate PDX models were then treated with Indo A (20 mg/kg) and tumor progression was assessed (Fig. 4h-i). Growth of the biomarker positive LCNEC model was significantly reduced whereas growth of the biomarker negative Merkel 2 model was not. These data support the ability of these biomarkers to predict responsiveness to anti-Cdk5 therapy.

## Enhanced drug delivery method

Although transgenic MTC mouse model animals responded favorably to Indo A treatments, tumor regression was not observed. Efforts to test higher doses of Indo A for increased potency were not feasible as cohorts receiving 30 mg/kg BW experienced 50% mortality while both 20 and 30 mg/kg treated cohorts evidenced some level of liver or kidney toxicity (Fig. 4n-o). To deliver Indo A selectively to tumors and thereby circumvent toxicity issues, we tested a biomimetic nanoparticle-based drug delivery system, leukosomes (LKs), generated from a combination of synthetic phospholipids and leukocyte membrane extracts^30,31^. Traditional nanoparticle delivery systems are dependent upon passive targeting through enhanced permeability and retention (EPR) of unhealthy tumor vasculature^32^. Although still benefiting from EPR, LKs actively target activated endothelium via mechanisms analogous to those utilized by white blood cells^33^. Importantly, LKs are successfully camouflaged by leukocyte membrane proteins leading to lower total opsonization and avoidance of the rapid immune clearance observed with purely synthetic nanoparticle platforms^34–36^.

As tumors are generally highly inflamed, we tested the ability of LKs to traffic to MTC tumors of NSE-p25OE mice and to stabilize lower doses of Indo A. Using intravital microscopy, LKs were verified to exhibit increased tumor localization compared to control liposomes and to time-dependently spread from the vasculature into surrounding tumor tissue (Fig. 4j-k). Maximum encapsulation of Indo A into LKs had no deleterious effects on particle size or homogeneity (Ext. Data Fig. 6) and allowed a dosage of 5 mg/kg BW Indo A per retro-orbital injection. HPLC/MS analysis of blood plasma demonstrated a 6-fold stabilization of encapsulated drug compared to free drug at these lower dosage levels (Fig. 4l).

Indo A delivered at 10 mg/kg BW in the free form had no effect on tumor growth in transgenic MTC animals (Fig. 4b). In contrast, delivering only 5 mg/kg BW Indo A encapsulated in LKs had the same effect as delivering 20 mg/kg BW free drug (Figs. 4b and 4m). Thus, the LK delivery system generated an equivalent effect utilizing a 75% lower dosage. LKs also protected animals against hepatic and renal toxicity (Fig. 4n-o). While complete tumor arrest or shrinkage was not observed at 5 mg/kg BW, further modifications to allow higher encapsulation of drug could provide additional benefit for this unique delivery approach.

## Discussion

The development of advanced sequencing techniques has led to an explosion of information pertaining to the genomic landscape of cancer. In some cases, this information successfully progressed to the development of personalized medicine. For example, mutations at Val600 in the serine/threonine kinase B-Raf are predictive of response to B-Raf inhibitors in patients with metastaic melanoma^37,38^. However, the majority of B-Raf mutant papillary thyroid cancer and colon cancer patients do not respond to B-Raf targeted inhibitors^39^, emphasizing the difficulty designing treatment options based on single gene mutations.

For NE neoplasms, mutations are prominent in genes encoding the scaffolding protein menin and the receptor tyrosine kinase RET. Menin mutations were initially identified over 20 years ago in patients with multiple endocrine neoplasia (MEN) type 1 syndrome^40,41^. Menin is a broadly expressed tumor suppressor in which mutations typically cause protein truncation^42,43^. Currently there are no therapeutics with the potential to circumvent these mutations.

Mutations in the proto-oncogene RET were discovered over 20 years ago in patients with MEN2 syndrome^44–46^. The development of vandetanib and carbozantinib, tyrosine kinase inhibitors that target RET, generated hope that patients possessing RET mutations could be successfully treated. Unfortunately, human trials revealed no correlation between the presence of a RET mutation and patient response to therapy^47–49^.

As with all cancers, many mutations, in addition to RET, are present within each MTC cancer cell. These additional mutations can alter the diverse input nodes of the signaling network that drive cancer cell growth and survival. For these reasons, looking at the signaling network with a broader lens that includes post-translational modifications could be beneficial and aid in the elimination of “false positive” non-responders that would be predicted responders from genomic or proteomic information alone. By assessing signaling states across a network of pathways, such an approach might also allow for more accurate stage-dependent therapeutic assessments to be made for individual patients. The current study reveals that Cdk5 is likely a contributor to at least a portion all NE tumor types. It also identifies a set of phosphorylation-based biomarkers which indicate that not only are Cdk5 pathway components present, but Cdk5 is actively modulating the signaling network and regulating cancer physiology.

In addition to being biomarkers of Cdk5 pathway activity, the phosphoproteins identified in this study are potentially directly involved in promotion of cell growth and/or survival. For example, RBL1 is a member of the retinoblastoma (RB) family of proteins that includes the tumor suppressor RB and RBL2. The RB family plays a major role in cell cycle regulation and is also involved in modulating senescence, apoptosis, and chromosomal stability^27^. Although functional compensation has been observed among the family members, some differences exist. Unlike RB, RBL1 and RBL2 do not bind to activating transcription factors E2F1-3. They interact instead with transcriptional repressors E2F4 and E2F5^50–52^. Remarkably, both RBL1 and RBL2, but not RB, are members of the DREAM complex, a regulatory unit that mediates cell entry into quiescence^53,54^.

Emerging information suggests Cdk5 may play a role in non-NE cancer as well^11^. As the heterogeneity between tumors of the same cancer type is becoming more apparent, the likelihood that treatment approaches will evolve based on individual tumor signaling states instead of general tumor type classification is increasing. The biomarkers identified in this study are not limited to use for NE cancer patients. Indeed, the Ovarian and PDAC PDX models analyzed here, both non-NE, exhibit high levels of NE features as well as biomarkers of Cdk5 pathway activation. A broader survey of PDX models for these novel biomarkers coupled with pre-clinical Cdk5 inhibitor testing could delineate finite cut-offs for classification of multiple forms of cancer as predicted responders for Cdk5-targeted therapy.

## Methods

### Animal Research

All animal work was performed in accordance with the guidelines of the Animal Welfare Act and the Guide for the Care and Use of Laboratory Animals under approved protocols by UTSW, UAB, and HMRI Institutional Animal Care and Use Committees.

### Antibody production and purification

Phosphopeptides (SIT*SPNRTGC-ASXL2, CAPSKLW*TPIKH-FAM53C, CSY*TPVKAIK-FLNB, CAPVEK*SPAK-H1B, CANYE*SPGKI-KNL2, CALA*TPQKNG-LARP6, CSIYI*SPHKN-RBL1, CAGLPG*SPKK-SUV39H1; *indicates a phosphorylated residue) were conjugated to *Limulus polyphemus* hemocyanin (Sigma H1757), emulsified with Freund’s adjuvant (Sigma F5881 or F5506), and injected subcutaneously into New Zealand White rabbits (Charles River Laboratories). Rabbits were boosted once and blood collected twice over a 5 week period for 12 months. Blood was allowed to clot at 4°C for 24 h, centrifuged at 1000 g, and plasma isolated and stored at −20°C.

Phosphorylation state-specific antibodies were purified from plasma using phosphopeptide affinity columns by elution with 100 mM Glycine pH 2.5 into 1 M Tris pH 8.6 (11:1 volume ratio, final 80 mM Tris pH ~7.5). Antibodies were dialyzed into 50 mM Tris pH 7.6 plus 150 mM NaCl and stored at −20°C.

### Cell culture and assays

All cells were tested and verified to be free of mycoplasma contamination. Cell lines were probed for NE markers to verify identity. Cells were cultured at 37°C and 5% CO_2_ in a humidified incubator. Fibroblasts were cultured in DMEM with 10% FBS. TT, MTC-SK, and SIN-J cells were cultured in Ham’s F12:Medium 199 (1:1) with 10% FBS. hPheo1, H146, and H1184 cells were cultured in RPMI with 10% FBS, 1 mM Na-pyruvate, and 10 mM HEPES. BON cells were cultured in DMEM: Ham’s F12 (1:1) with 10% FBS. INS cells were cultured in RPMI with 10% FBS, 1 mM Na-pyruvate, 10 mM HEPES, 4.5 g/L glucose, and 50 μM β-ME. TT-RLuc cells were cultured in RPMI with 20% FBS, 100 μg/ml penicillin, and 100 μg/ml streptomycin. *Growth assays.* Cells were seeded onto a black 96-well plates with a clear optical bottom and allowed to recover overnight. Growth of hPheo1 cells was measured 2 days after inhibitor treatment and growth of Fibro, TT, MTC-SK, BON, INS, and H146 cells was measured 6 days after inhibitor treatment using Cyquant Direct Proliferation Assay (Invitrogen) and an Optima Fluostar plate reader (BMG LabTech). For SIP experiments, cells received two SIP treatments per experiment. Growth of BON and hPheo1 cells was measured 2 days after initial SIP treatment and growth of Fibro and MTC-SK cells was measured 6 days after initial SIP treatment using procedures described above.

#### Immunoblot analysis

Cells were seeded onto 6 well dishes and allowed to recover overnight. Cells were treated for 4 h with inhibitors and then lysed in 50 mM NaF and 1% SDS with brief sonication. Samples were diluted to equivalent total protein concentrations in 1X Laemmli buffer and separated by SDS-PAGE. Proteins were transferred onto nitrocellulose for immunoblotting utilizing in-house phosphorylation state-specific antibodies, anti-Cdk5 (sc-173), anti-p35 (sc-820), anti-GAPDH (Sigma G8795), anti-actin (Abcam ab6276), anti-ASXL2 (Abcam ab106540, Sigma sab1407639), anti-Fam53C (Abcam 105679), anti-FLNB (Abnova PAB30702), anti-H1B (sc-247158), anti-KNL2 (sc-162587), anti-RBL (sc-318-G), anti-LARP6 (Sigma sab1407657), and anti-SUV39H1 (Sigma S8316). Anti-goat, -rabbit, and -mouse secondary antibodies conjugated to either IRdye 680RD or IRdye 800CW (LiCor) were used for detection on a LiCor Odessey CLx. Actin and GAPDH were used as sample processing controls. Other total proteins were probed on the same membrane as phosphoproteins unless antibody conditions did not allow.

### Human tissue analysis

#### Collection

Human tissues were collected with patient consent and in accordance with institutional review board (IRB) regulations. Samples were collected under University of Wisconsin Madison IRB 2011-0145, MD Anderson IRB PA11-0744, University of Texas Southwestern Medical Center IRB STU102010-042 and STU102010-051, NICHD IRB 00-CH-0093, Louisiana State University IRB 5774, and University of Sydney LNR/13/HAWKE/424 – 1312-417M.

#### Histology

Formalin-fixed, paraffin-embedded samples were cut into 5 μm sections, deparaffinized, and subjected to microwave antigen retrieval (citrate buffer, pH 6.0). Sections were then stained using standard protocols for H&E or immunostained with antibodies recognizing p35/p25 (sc-820, Santa Cruz Biotechnology) or Cdk5 (308-Cdk5, PhosphoSolutions). For immunostaining, sections were permeabilized with 0.3% Triton X-100, quenched free of endogenous peroxidases, and blocked with 2.5% normal goat serum prior to overnight incubation with primary antibodies at 4°C. Bound primaries were detected by sequential incubation with biotinylated-secondary antisera, streptavidin-peroxidase (Vector Laboratories), and diaminobenzidine chromagen (DAKO) following the manufacturer’s directions.

#### Immunoblot analysis

Tissues were crushed while frozen then lysed in 50 mM NaF and 1% SDS with brief sonication. Samples were diluted to equivalent total protein concentrations in 1X Laemmli buffer and separated by SDS-PAGE. Proteins were transferred onto nitrocellulose for immunoblotting as described above.

### Intravital Microscopy

IVM was performed using an upright Nikon A1R laser scanning confocal microscope with a resonance scanner, motorized and heated stage, and Nikon long-working distance 4× and 20× dry plan-apochromat objectives housed within the IVM Core at the HMRI. For imaging, NSE-p25OE mice were anesthetized with isoflurane and the ventral surface of the neck opened to expose the trachea, salivary glands, and MTC tumors. Tumors were positioned in direct contact with the coverslip, visualized using the GFP signal, and positions selected for imaging. After selection of positions, Cy5.5 labeled LKs and LPs were administered via retro-orbital injection and mice were imaged continuously for 2 hours using the 4x objective. Images were quantified using Nikon Elements. The tumor accumulation reported was normalized by dividing the area occupied by LKs or LPs by the area occupied by the tumor within each image.

### LCMS2 Analysis of IndoA

#### Extraction

Sera (25 μL) were reconstituted in 25 μL of extraction solution (25% acetonitrile/75% H_2_0), vortexed for 1 min, diluted with 25 μL of 100% ACN, followed by two cycles of vortexing for 1 min and incubating at room temperature (RT) for 10 min. Samples were then diluted with 200 μL of 100% ACN, vortexed for 1 min, then stored at −20°C overnight. Samples were thawed at RT for ~10 min, centrifuged at 14,000 g for 10 min at 4°C to remove cell debris, then supernatants were transferred to 2 mL tinted glass vials and dried down to ~5-10 μL under argon gas at 25°C for 25 min. Samples were immediately reconstituted (adjusted to 210 μL) in 25% ACN/1% trifluoroacetic acid/74% H_2_0 and centrifuged at 14,000 g for 10 min at 4°C to remove any residual debris. The supernatants were then transferred to tinted vials prior to analysis.

#### LC/MS

Analysis was performed on a Dionex Ultimate 3000 UHPLC+ Focused Stack & Auto Sampler (Thermo Fisher Scientific/Dionex) a using a RP C18 Hypersil Gold (100mm I.D. x 4.6mm, 5 mm 175Å pore size; Thermo Fisher Scientific) in-line with an LTQXL mass spectrometer equipped with a heated electrospray ionization source (Thermo Fisher Scientific), and all data were collected in selective reaction monitoring (SRM) mode. The HPLC was configured with binary mobile phases that include solvent A (0.1% TFA/99.9% H_2_O), and solvent B (0.1% TFA/15% H_2_O/5% ACN). The gradient program steps were run in linear mode as follows; 0-6 min at 75%-50% B (200 μL/min), 6-7 min at 50%-80% B (200 μL/min), 7-11min at 80%-90% B (200 μL/min), 11-12min at 90%-25% B (500 μL/min), and finally 12-16min at 25%B (200 μL/min). SRM mode was optimized using a parent ion window of 453.2 +/− 1.0 *m/z*, 20% normalized collision energy, activation energy at 0.240, activation time of 30 ms, with a daughter ion window of 306.5 +/− 1.5 *m/z*. The resultant Xcalibur RAW files were collected in profile mode and the SRM base peak values processed and extracted using Xcalibur version 2.2 sp1.48.

### Leukosome synthesis and characterization

LKs were developed as previously reported^31^. Briefly, 1,2- dipalmitoyl-sn-glycero-3-phosphocholine (DPPC) and 1,2-dioleoylsn-glycero-3-phosphocholine (DOPC) and cholesterol (Avanti Polar Lipids) (4:3:3 molar ratio) were dissolved in ethanol at a final lipid concentration of 9 mM and mixed with membrane proteins, previously resuspended in aqueous buffer at 1:50 protein to-lipid concentrations, using the NanoAssemblr Benchtop platform (Precision NanoSystems, Inc.). Passive loading of Indo A within LKs was obtained by dissolving the drug in the ethanol mixture containing the lipids. Size and polydispersity index were determined through dynamic light scattering analysis using a Nanosizer ZS (Malvern Instruments). Surface charge (Zeta potential) was measured using a ZetaSizer Nano ZS (Malvern Instruments).

### Magnetic Resonance Imaging

MRI conducted at UT Southwestern was performed using a 7-Tesla small animal MRI system (Agilent Inc.) with a 40 mm (i.d.) radio frequency (RF) coil and a 400mT/m gradient coil set. Animals were anesthetized by isoflurane and imaged in a supine position, head first with the thyroid centered with respect to the center of a RF coil. Two-dimensional (2D) fast spin-echo (FSE) images on three orthogonal planes (transverse, coronal and sagittal) were firstly acquired to ensure the position and the orientation of the thyroid tumors. For volume measurements of the thyroid tumors, the high resolution T2-weighted FSE axial images was acquired. Major imaging parameters were: TR/TE = 2500/40 msec, FOV = 25.6 x 25.6 mm, matrix size = 256 x 256, slice thickness = 1 mm, no gap, 8 averages, affording 100 μm inplane resolution.

### Phosphoproteomics

#### Peptide preparation

Mouse tumors were homogenized in 8 M urea lysis buffer (20 mM HEPES pH 8.0, 9 M urea, 1 mM sodium vanadate, 2.5 mM sodium pyrophosphate, 1 mM ß-glycerol-phosphate), sonicated, then centrifuged for 15 min at 4°C at 20,000 g. Supernatants were reduced with 4.5 mM DTT for 30 min at 55°C followed by alkylation with 10 mM iodoacetamide. The samples were then digested with trypsin overnight at room temperature. Digests were acidified with 1% TFA and peptides desalted and purified over Sep-Pak C18 columns (Waters, WAT051910) using 40% acetonitrile in 0.1% TFA for elution. Elutes were lyophilized and stored at −80°C.

#### Immunoaffinity purification (IAP) of peptides

Lyophilized peptides were dissolved in IAP buffer (50 mM MOPS pH 7.2, 10 mM sodium phosphate, 50 mM NaCl), sonicated, and insoluble matter removed by centrifugation. CDK substrate motif antibody (Cell Signaling Technology #2324) and MAPK substrate motif antibody (Cell Signaling Technology #2325) was coupled to protein A beads (Roche). Immobilized antibody was incubated with peptide mixtures, immunoprecipitation was carried out at 4 ° C overnight, then washed with IAP buffer, and eluted with 0.15% TFA. Eluates were further purified using C18 microtips (StageTips or Eppendorf C18 PerfectPure tips) with elution in 60% MeCN, 0.1% TFA then lyophilized.

#### Analysis by LC-MS/MS

Purified peptide mixtures were loaded onto a 10 cm x 75 μm PicoFrit capillary column (New Objective) packed with Magic C18 AQ reversed-phase resin (Michrom Bioresources) using a Famos autosampler with an inert sample injection valve (Dionex). The column was developed with a 45-min gradient of acetonitrile in 0.125 % formic acid (Ultimate pump, Dionex), and tandem mass spectra were collected in a data-dependent manner with a Thermo Fisher Scientific LTQ ion trap mass spectrometer equipped with Electron Transfer Dissociation (ETD) module or with an Orbitrap mass spectrometer.

#### Assigning peptide sequences using Sequest

MS/MS spectra were evaluated using TurboSequest in the Sequest Browser package supplied as part of BioWorks 3.3 (Thermo Fisher Scientific). Searches were performed against the NCBI human protein database. Cysteine carboxamidomethylation was specified as a static modification and phosphorylation was allowed as a variable modification on serine and threonine.

### Preclinical Drug Testing

A bi-transgenic mouse model of MTC was generated as previously described by crossing NSE-rTA and TetOp-p25GFP parental lines^12^. Bi-transgenic litters were monitored by MRI. Beginning at 10-40 mm^3^ bilateral tumor volume, animals were treated once every 3 days by IP with vehicle (0.7% DMSO, 3.4% EtOH, 7.4% PEG400, 3.4% PG, 3.4% Kolliphor EL, 1.1% Tween 80 in 1X PBS), 10 mg/kg, 20 mg/kg, or 30 mg/kg BW Indo A. Animals were monitored by MRI for 2 weeks then sacrificed 24 h post-final injection by CO_2_ euthanasia and cardiac perfusion with 0.1 mM Ammonium molybdate, 5 mM EGTA, 50 mM NaF, 2 mM Na orthovanidate, 10 mM Na pyrophosphate, and protease inhibitors (Sigma S8820) in PBS. Tissues were frozen or fixed in 4% PFA for 24 h and submitted for paraffin embedding. Human BON and TT cell xenograft mouse models were generated by implanting 5e^6^ BON or TT-RLuc cells subcutaneously in the NU/NU Nude Mouse (Crl:NU-Foxn1^nu^) strain from Jackson laboratories. Beginning at tumor volumes of 100-450 mm^3^ (average starting sizes between groups varied less than 40mm^3^), animals were treated as described above and monitored by caliper measurement for 2 weeks. Animals were sacrificed and tissue processed as described above. All animals were randomly assigned to treatment groups but blinding was not possible.

### Scanning Electron Microscopy

Cells were fixed on coverslips with 2.5% (v/v) glutaraldehyde in 0.1 M sodium cacodylate buffer overnight at 4°C. After three rinses in 0.1 M sodium cacodylate buffer, they were post-fixed with 1% osmium tetroxide in 0.1 M sodium cacodylate buffer for 45 min. Cells were rinsed with water and dehydrated with increasing concentration of ethanol, followed by increasing concentrations of Hexamethyldisilazane in ethanol. Cells were air dried under the hood, mounted on SEM stubs and sputter coated with gold palladium in a Cressington 108 auto sputter coater. Images were acquired on a Field-Emission Scanning Electron Microscope (Zeiss Sigma) at 3.00 kV accelerating voltage.

### Statistical Analysis

Pre-clinical drug testing in animal models and accumulation of nanoparticles were analyzed by two-way ANOVA with repeated measures. All indications of significance are for the treatment group over the entire time period, not individual time points. Immunoblots containing >2 conditions, AST assays and Urea assays were analyzed by one-way ANOVA. Correlations in human tumor data were performed by Spearman rank-order analysis. Simple comparisons of two groups were performed using twotailed Student’s t-test. Sample sizes are provided with figure legends or in results. (*p<0.05, **p<0.01, ***p<0.005, ****p<0.001).

### Toxicity Assays

Prior to cardiac perfusion, blood was collected from animal subjects via retro-orbital bleeding. Blood was immediately mixed with EDTA (10 mM final) then centrifuged at 1000 g for 10 min at 4°C to allow isolation of plasma. AST assays (Sigma MAK055) and Urea assays (Sigma MAK006) were performed on plasma per the manufacturer’s instructions.

We thank Melanie Cobb, Joseph Goldstein, John Minna, Roswitha Pfragner, and Courtney Townsend for cell lines and Eric Knudsen for primary fibroblast cultures. We thank Haydn Ball and the UTSW Protein Technology Core for peptide synthesis, the UTSW Animal Resource Center for antigen injection and blood collection, Robyn Leidel and Kate Luby-Phelps for help with SEM, Kelly Hartman for assistance with leukosome preparation, John Totenhagen and Samuria Thomas and the UAB Small Animal Imaging Facility for additional MRI, and HMRI Intravital Microscopy Core for IVM. We thank Champions Oncology for PDX tissue and preclinical testing in PDX models, and Pfizer for CP681301.

This work was supported by an American Cancer Society Postdoctoral Fellowship (A.M.C) and the Sackler Foundation (A.M.C. and J.A.B.); NIH award S10OD023552-01 (M.T.); an American Thyroid Association Research Grant and the Dedman Family Scholar in Clinical Care Award (S.C.O.); an American Cancer Society Institutional Research Grant Junior Faculty Development Award (S.R.) and the SDHB PHEO/Para Coalition (S.R. and J.A.B.); the William Randolph Heart Foundation, the Robert J. Kleberg, Jr. and Helen C. Kleberg Foundation, Cancer Prevention & Research Institute of Texas Award ID RP170466, and NCI and the Office of Research on Women’s Health Grant # 1R56CA213859 (E.T.); an American Cancer Society Research Scholars Award, and NIH awards DA033485-01, MH083711-01, NS073855-01, and R56MH116896 (J.A.B); and NCI awards 5P30CA142543 (UTSW Simmons Comprehensive Cancer Center) and P30CA013148 (UAB O’Neal Comprehensive Cancer Center).

A.M.C. and J.A.B. conceptualized the study. A.M.C. performed cell culture assays, analysis of phosphoproteomic data, production of phosphorylation state-specific antibodies, immunoblotting, enzyme and urea assays, quantitation of MRI, IVIS imaging, and contributed to generation of mouse xenograft models, drug treatments in mouse models, Indo A biodistribution analysis, and performed data analysis. C.T. performed IHC. K. Pozo contributed to phosphoproteomics and SEM. R.T. contributed to work in mouse models. R.M. designed, synthesized, and characterized leukosomes. A.G. and T.W. performed phosphoproteomics. E.D.R. and J.O.M. performed IVM. S.Z. and M.T. performed small animal MRI. F.G. and Boehringer Ingelheim provided Indo A and Indo B. R.J. contributed to generation of xenograft mouse models. J.A.M. performed LC-MS for Indo A biodistribution analysis. N.K. and M.S.M. performed IPA analysis. H.K.G. provided the hPheo1 cell line. S.C.O., F.E.N., and S.R. contributed financially. H.C., E.G.G., K.J.H., K. Pacak, A.J.G., B.R., and E.A.W. provided human biospecimens. J.A.B., E.T., and H.C. supervised the study and data interpretation. A.M.C. and J.A.B. wrote the manuscript. All authors reviewed and edited the manuscript.

Phosphoproteomic data are deposited in PhosphoSitePlus (www.phosphositeplus.org). Reprints and permissions information is available at www.nature.com/reprints. The authors declare no competing financial interests. Readers are welcome to comment on the online version of the paper. Correspondence and requests for materials should be addressed to J.A.B.

## Extended Data

**Extended Data Fig. 1:**
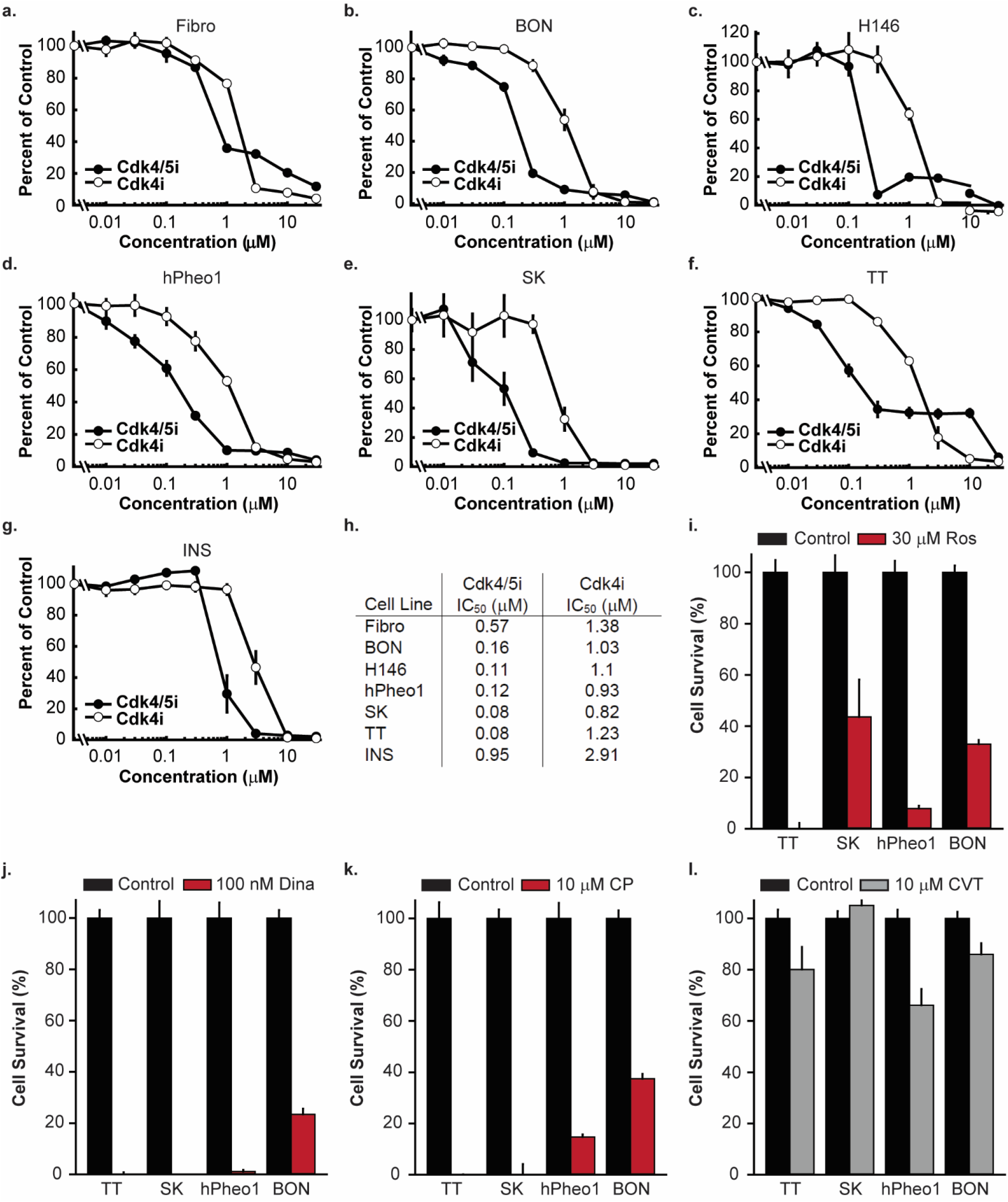
Cdk5 inhibition blocks growth of NE cancer cells. **a-g**, Cells were treated with increasing concentrations of Indo A (Cdk4/5i; n=6-9) and Indo B (Cdk4i; n=6-9) and monitored for effects on cell growth. **h**, IC_50_ values for each cell line and inhibitor were calculated by 4-parameter logistic regression. **i-l**, NE cell lines were treated with control or the Cdk inhibitor indicated; Roscovitine (Ros; n=4-8), Dinacyclib (Dina; n=6-8), CP681301 (CP; n=4-8), or CVT313 (CVT; n=8-12).

**Extended Data Fig. 2:**
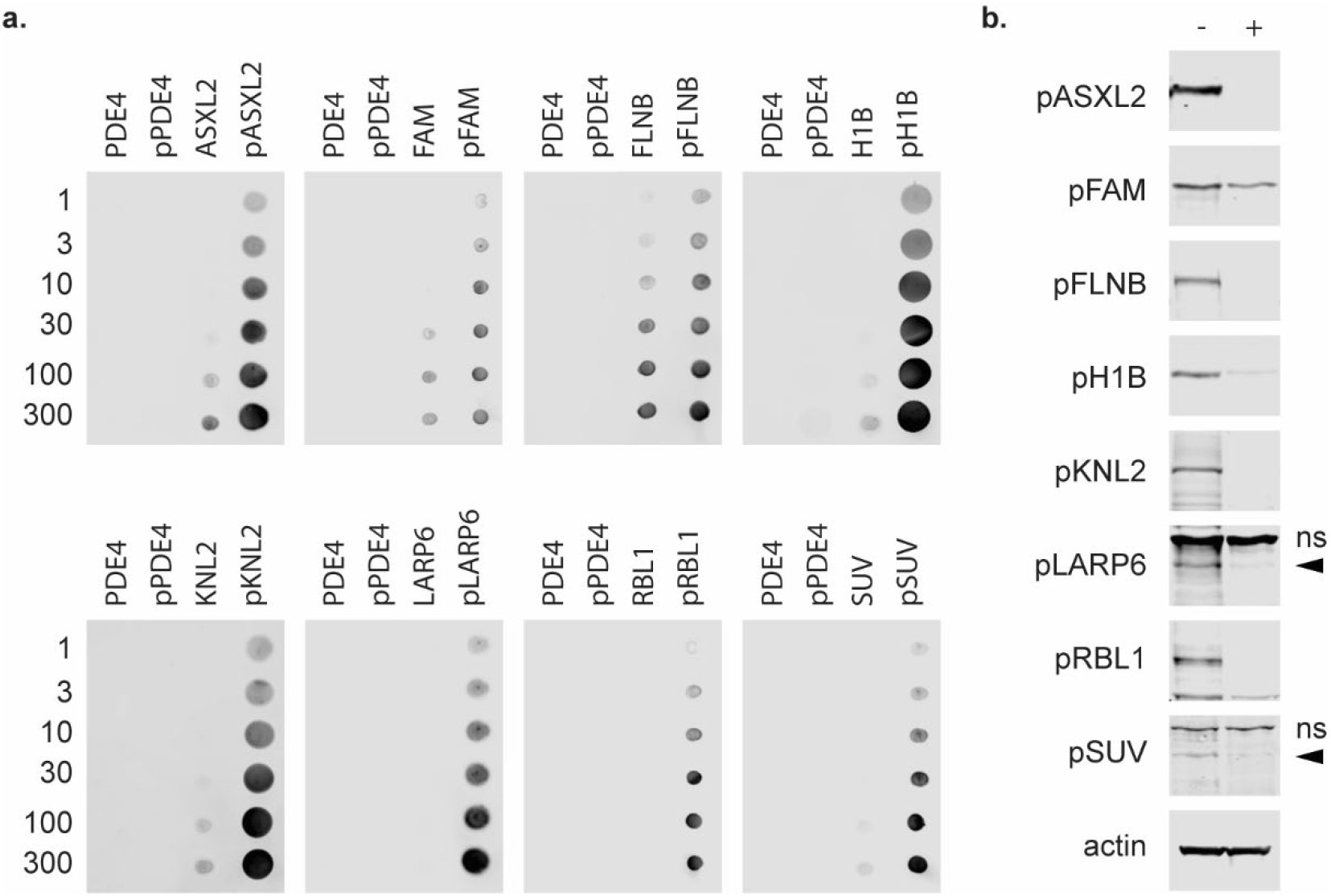
Phosphorylation state-specific antibody validation. **a**, Dot blots of nitrocellulose spotted with increasing pmol of PDE4 peptide, phospho-PDE4 (pPDE4) peptide, peptide of interest, or phospho-peptide of interest and probed with affinity purified anti-sera. **b**, Immunoblot analysis of NSE-p25OE tumor lysate, treated with (+) and without (-) lambda protein phosphatase, probed with affinity purified anti-sera. Non-specific bands (ns), specific bands denoted by arrowheads where applicable.

**Extended Data Fig. 3:**
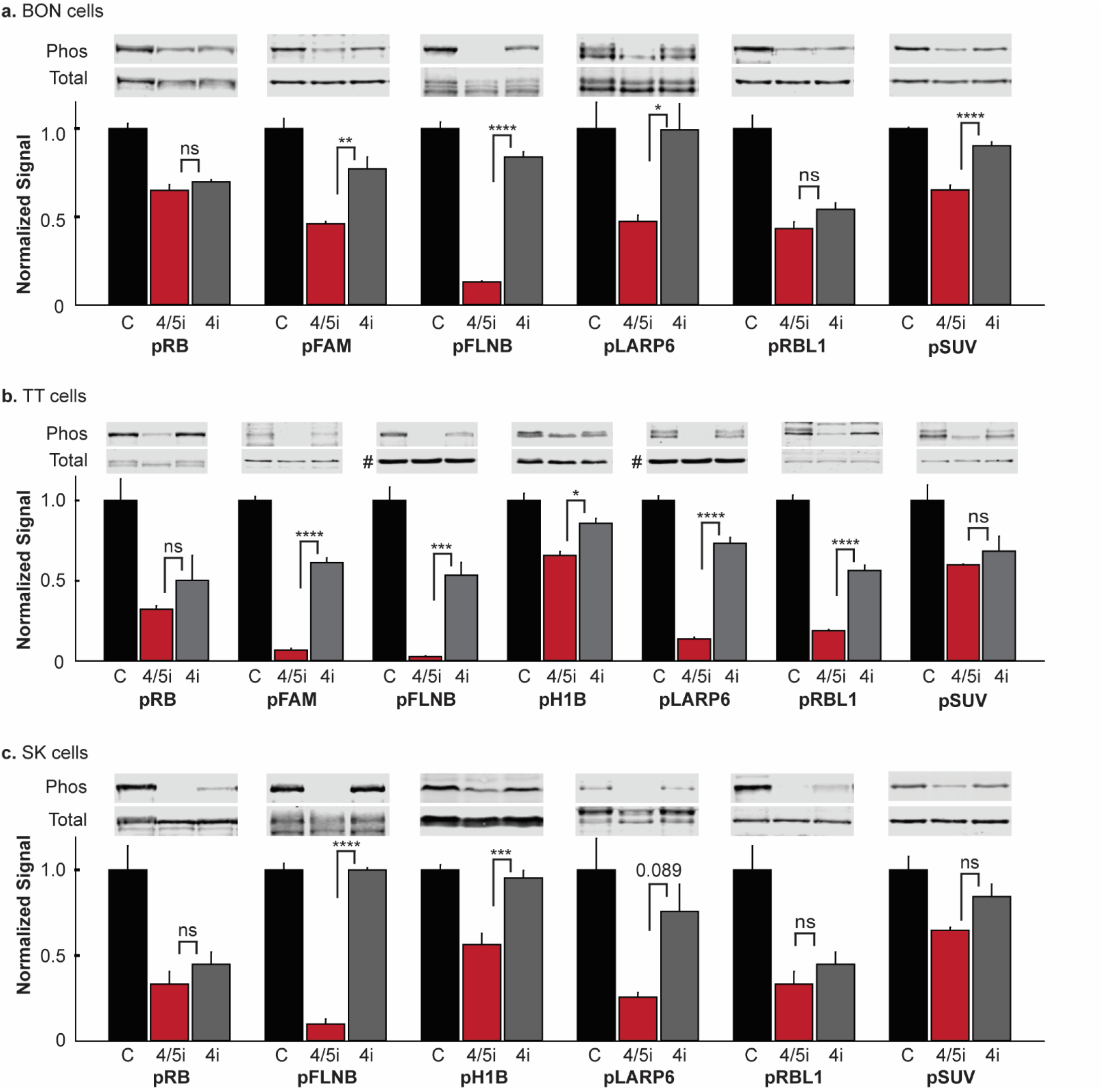
Phosphoproteins are dependent on Cdk5 activity. **a-c**, Immunoblot analysis of phosphoproteins in BON cells (**a**), TT cells (**b**), and MTC-SK cells (**c**) treated with 0.3% DMSO (Control), 2 μM Indo A (Cdk4/5i), or 2 μM Indo B (Cdk4i) for 4 h. For **b**, pFLNB and pLARP are normalized to actin (#); all others are normalized to each specific protein. Statistics by One-way ANOVA, n=3.

**Extended Data Fig. 4:**
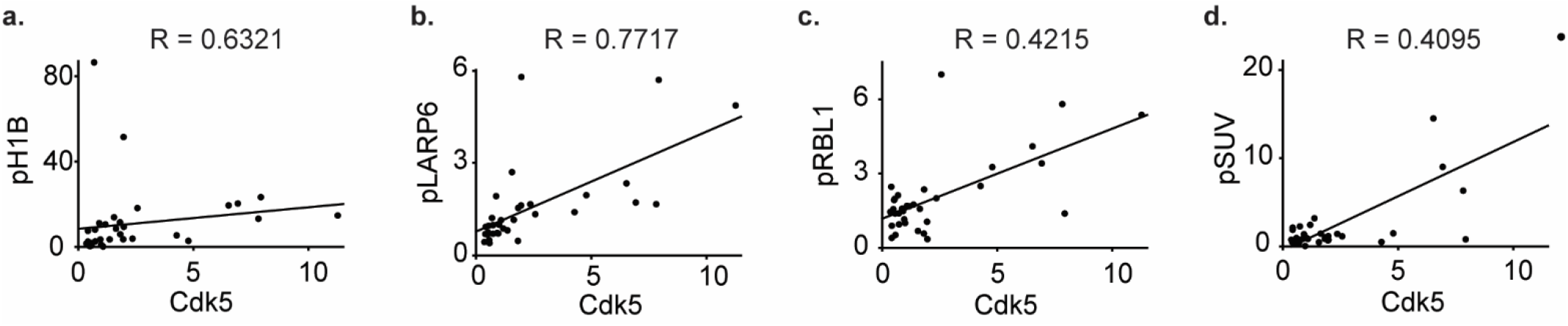
Phosphoprotein levels correlate with Cdk5 expression. Phosphoprotein level versus Cdk5 expression in samples represented in Fig. 3c-d analyzed by Spearman rank-order analysis; rho (R).

**Figure 5:**
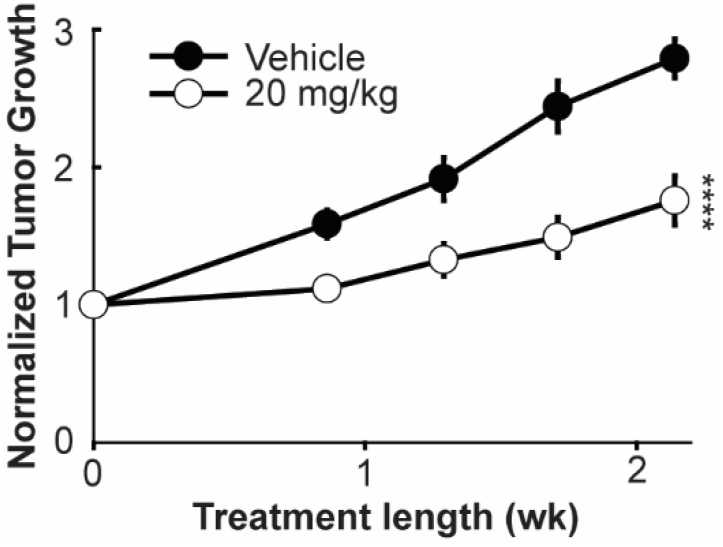
Inhibition of Cdk5 activity suppresses growth of pancreatic NE tumors. Quantitation of tumor growth over time from caliper measurements of BON xenograft pancreatic NE tumor model mice treated with vehicle or 20 mg/kg BW Indo A. (n=8-9).

**Figure 6:**
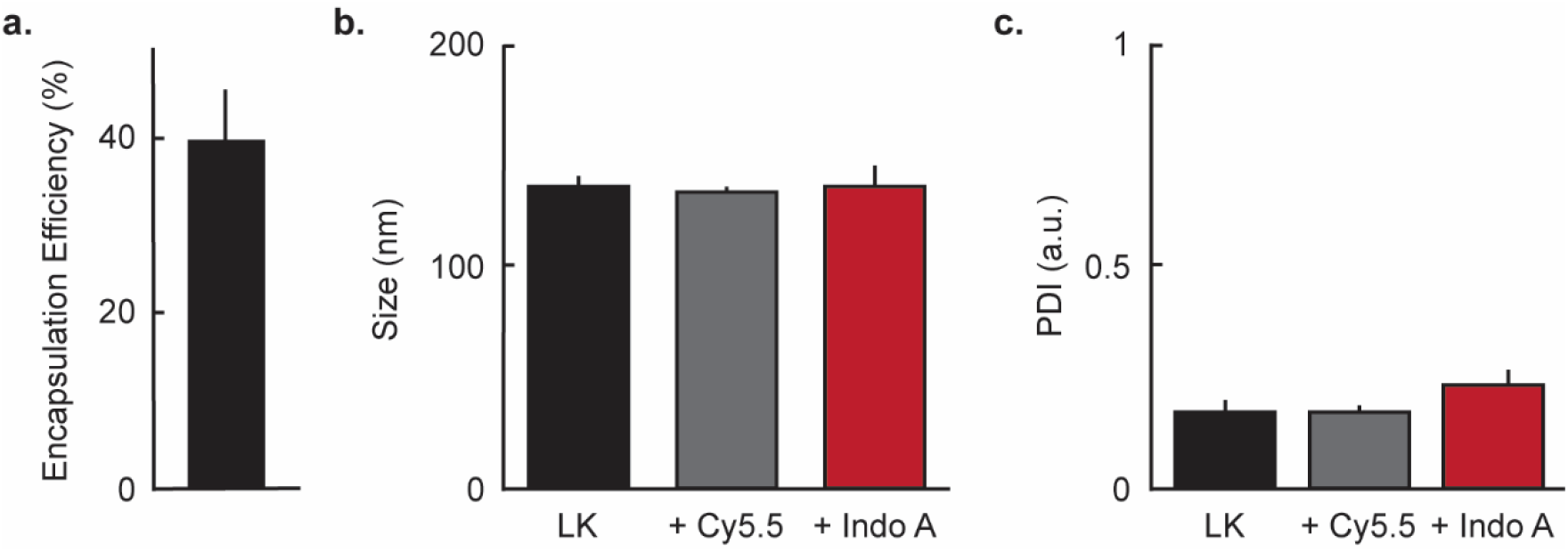
Encapsulation of Indo A does not affect biophysical properties of LKs. **a**, Entrapment efficiency of Indo A into LKs as determined by intrinsic Indo A fluorescence. **b-c**, Dynamic light scattering analysis of LKs, Cy5.5 labeled LKs, and Indo A loaded LKs to determine size (**b**) and polydispersity (**c**).

**Extended Data Table 1:**
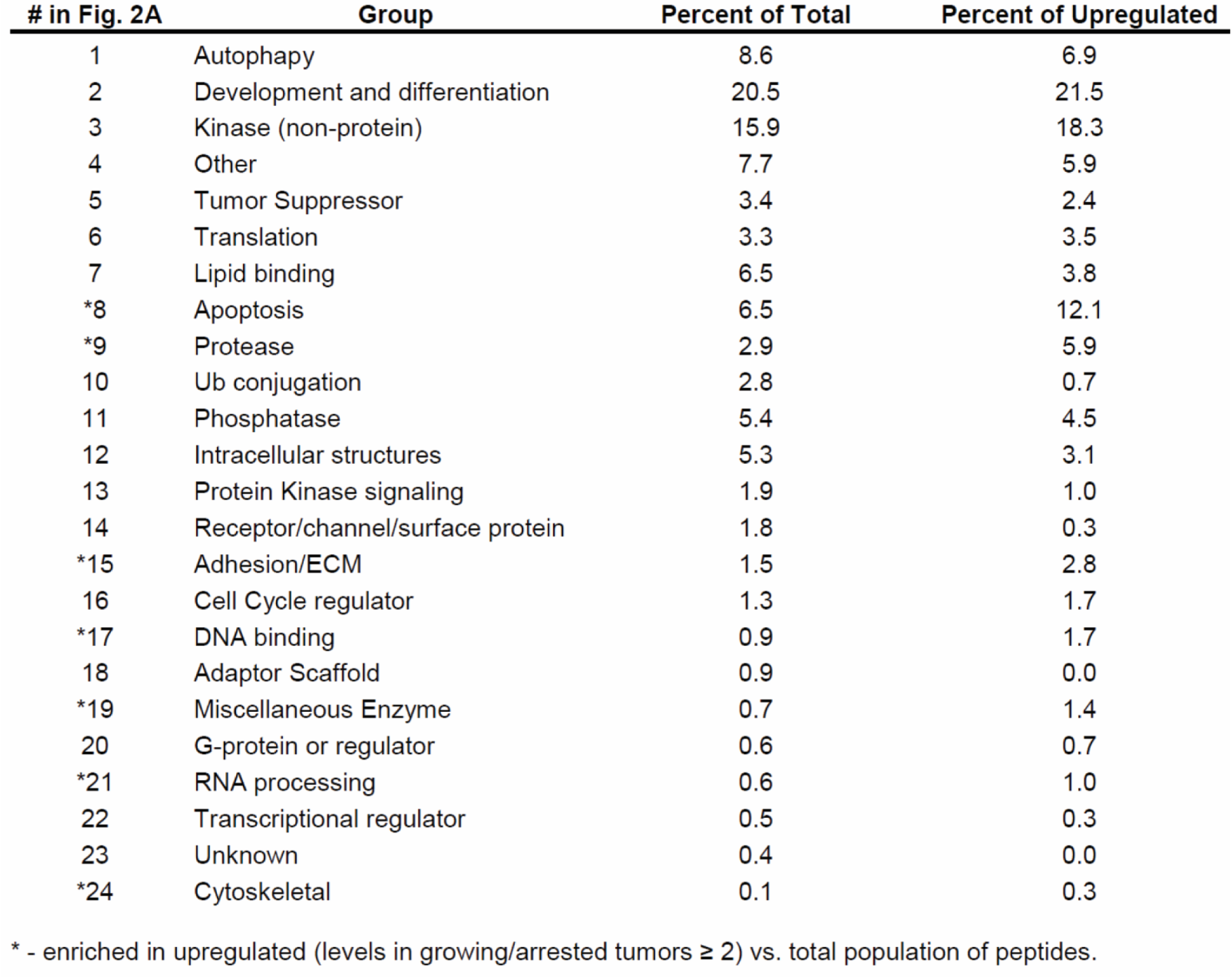
Protein groups of peptides identified in LC-MS/MS of MTC mouse tumors.

| # in Fig. 2A | Group | Percent of Total | Percent of Upregulated |
| --- | --- | --- | --- |
| 1 | Autophagy | 8.6 | 6.9 |
| 2 | Development and differentiation | 20.5 | 21.5 |
| 3 | Kinase (non-protein) | 15.9 | 18.3 |
| 4 | Other | 7.7 | 5.9 |
| 5 | Tumor Suppressor | 3.4 | 2.4 |
| 6 | Translation | 3.3 | 3.5 |
| 7 | Lipid binding | 6.5 | 3.8 |
| *8 | Apoptosis | 6.5 | 12.1 |
| *9 | Protease | 2.9 | 5.9 |
| 10 | Ub conjugation | 2.8 | 0.7 |
| 11 | Phosphatase | 5.4 | 4.5 |
| 12 | Intracellular structures | 5.3 | 3.1 |
| 13 | Protein Kinase signaling | 1.9 | 1.0 |
| 14 | Receptor/channel/surface protein | 1.8 | 0.3 |
| *15 | Adhesion/ECM | 1.5 | 2.8 |
| 16 | Cell Cycle regulator | 1.3 | 1.7 |
| *17 | DNA binding | 0.9 | 1.7 |
| 18 | Adaptor Scaffold | 0.9 | 0.0 |
| *19 | Miscellaneous Enzyme | 0.7 | 1.4 |
| 20 | G-protein or regulator | 0.6 | 0.7 |
| *21 | RNA processing | 0.6 | 1.0 |
| 22 | Transcriptional regulator | 0.5 | 0.3 |
| 23 | Unknown | 0.4 | 0.0 |
| *24 | Cytoskeletal | 0.1 | 0.3 |

\* - enriched in upregulated (levels in growing/arrested tumors $\geq 2$ ) vs. total population of peptides.

**Extended Data Table 2:**
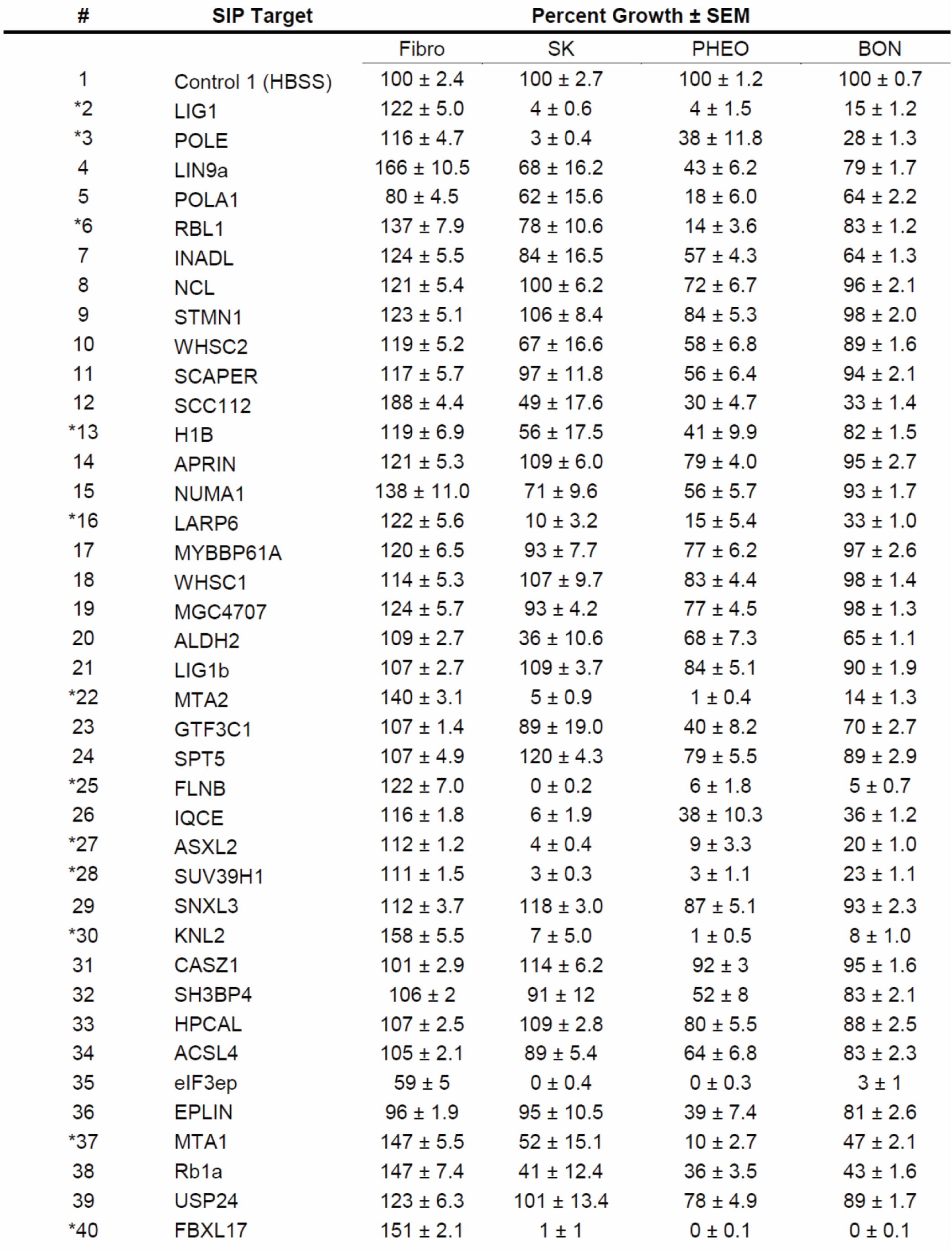
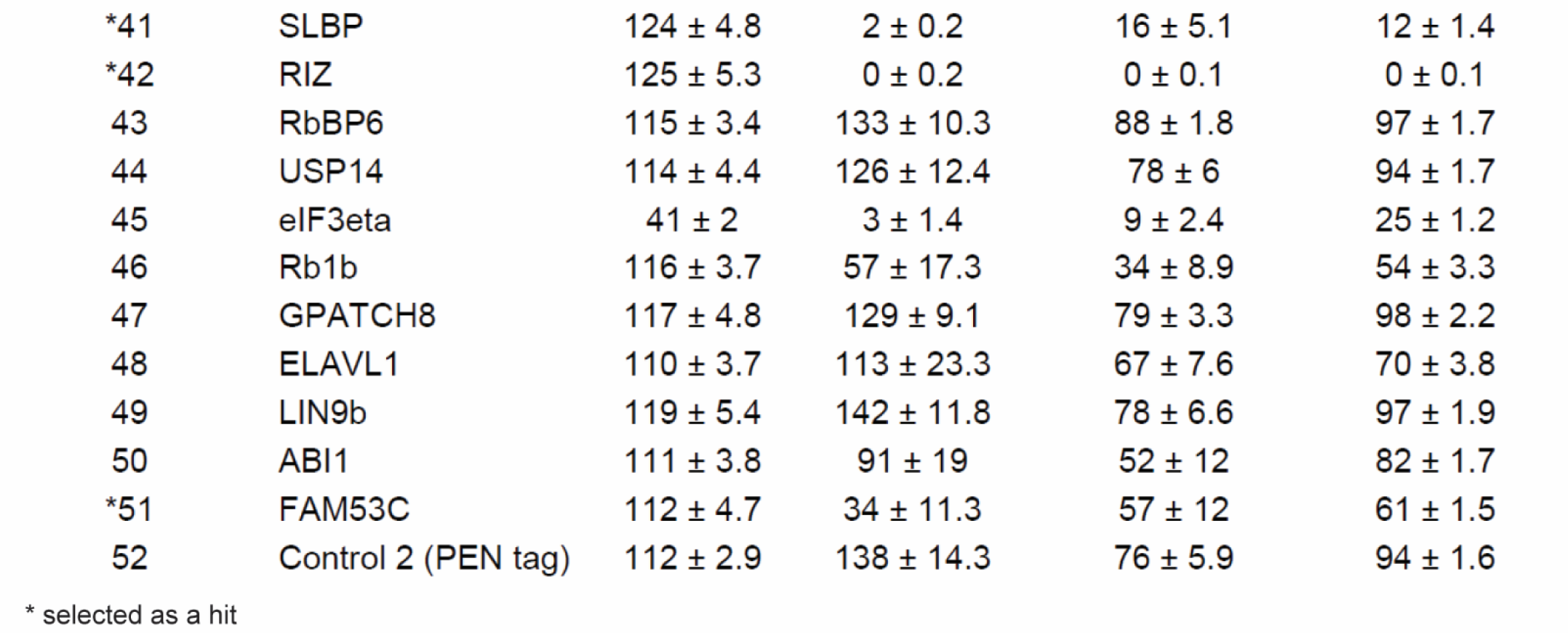
Growth inhibition by individual SIPs represented in Fig. 2b.

| # | SIP Target | Percent Growth ± SEM | | | |
| --- | --- | --- | --- | --- | --- |
|  |  | Fibro | SK | PHEO | BON |
| 1 | Control 1 (HBSS) | 100 ± 2.4 | 100 ± 2.7 | 100 ± 1.2 | 100 ± 0.7 |
| *2 | LIG1 | 122 ± 5.0 | 4 ± 0.6 | 4 ± 1.5 | 15 ± 1.2 |
| *3 | POLE | 116 ± 4.7 | 3 ± 0.4 | 38 ± 11.8 | 28 ± 1.3 |
| 4 | LIN9a | 166 ± 10.5 | 68 ± 16.2 | 43 ± 6.2 | 79 ± 1.7 |
| 5 | POLA1 | 80 ± 4.5 | 62 ± 15.6 | 18 ± 6.0 | 64 ± 2.2 |
| *6 | RBL1 | 137 ± 7.9 | 78 ± 10.6 | 14 ± 3.6 | 83 ± 1.2 |
| 7 | INADL | 124 ± 5.5 | 84 ± 16.5 | 57 ± 4.3 | 64 ± 1.3 |
| 8 | NCL | 121 ± 5.4 | 100 ± 6.2 | 72 ± 6.7 | 96 ± 2.1 |
| 9 | STMN1 | 123 ± 5.1 | 106 ± 8.4 | 84 ± 5.3 | 98 ± 2.0 |
| 10 | WHSC2 | 119 ± 5.2 | 67 ± 16.6 | 58 ± 6.8 | 89 ± 1.6 |
| 11 | SCAPER | 117 ± 5.7 | 97 ± 11.8 | 56 ± 6.4 | 94 ± 2.1 |
| 12 | SCC112 | 188 ± 4.4 | 49 ± 17.6 | 30 ± 4.7 | 33 ± 1.4 |
| *13 | H1B | 119 ± 6.9 | 56 ± 17.5 | 41 ± 9.9 | 82 ± 1.5 |
| 14 | APRIN | 121 ± 5.3 | 109 ± 6.0 | 79 ± 4.0 | 95 ± 2.7 |
| 15 | NUMA1 | 138 ± 11.0 | 71 ± 9.6 | 56 ± 5.7 | 93 ± 1.7 |
| *16 | LARP6 | 122 ± 5.6 | 10 ± 3.2 | 15 ± 5.4 | 33 ± 1.0 |
| 17 | MYBBP61A | 120 ± 6.5 | 93 ± 7.7 | 77 ± 6.2 | 97 ± 2.6 |
| 18 | WHSC1 | 114 ± 5.3 | 107 ± 9.7 | 83 ± 4.4 | 98 ± 1.4 |
| 19 | MGC4707 | 124 ± 5.7 | 93 ± 4.2 | 77 ± 4.5 | 98 ± 1.3 |
| 20 | ALDH2 | 109 ± 2.7 | 36 ± 10.6 | 68 ± 7.3 | 65 ± 1.1 |
| 21 | LIG1b | 107 ± 2.7 | 109 ± 3.7 | 84 ± 5.1 | 90 ± 1.9 |
| *22 | MTA2 | 140 ± 3.1 | 5 ± 0.9 | 1 ± 0.4 | 14 ± 1.3 |
| 23 | GTF3C1 | 107 ± 1.4 | 89 ± 19.0 | 40 ± 8.2 | 70 ± 2.7 |
| 24 | SPT5 | 107 ± 4.9 | 120 ± 4.3 | 79 ± 5.5 | 89 ± 2.9 |
| *25 | FLNB | 122 ± 7.0 | 0 ± 0.2 | 6 ± 1.8 | 5 ± 0.7 |
| 26 | IQCE | 116 ± 1.8 | 6 ± 1.9 | 38 ± 10.3 | 36 ± 1.2 |
| *27 | ASXL2 | 112 ± 1.2 | 4 ± 0.4 | 9 ± 3.3 | 20 ± 1.0 |
| *28 | SUV39H1 | 111 ± 1.5 | 3 ± 0.3 | 3 ± 1.1 | 23 ± 1.1 |
| 29 | SNXL3 | 112 ± 3.7 | 118 ± 3.0 | 87 ± 5.1 | 93 ± 2.3 |
| *30 | KNL2 | 158 ± 5.5 | 7 ± 5.0 | 1 ± 0.5 | 8 ± 1.0 |
| 31 | CASZ1 | 101 ± 2.9 | 114 ± 6.2 | 92 ± 3 | 95 ± 1.6 |
| 32 | SH3BP4 | 106 ± 2 | 91 ± 12 | 52 ± 8 | 83 ± 2.1 |
| 33 | HPCAL | 107 ± 2.5 | 109 ± 2.8 | 80 ± 5.5 | 88 ± 2.5 |
| 34 | ACSL4 | 105 ± 2.1 | 89 ± 5.4 | 64 ± 6.8 | 83 ± 2.3 |
| 35 | eIF3ep | 59 ± 5 | 0 ± 0.4 | 0 ± 0.3 | 3 ± 1 |
| 36 | EPLIN | 96 ± 1.9 | 95 ± 10.5 | 39 ± 7.4 | 81 ± 2.6 |
| *37 | MTA1 | 147 ± 5.5 | 52 ± 15.1 | 10 ± 2.7 | 47 ± 2.1 |
| 38 | Rb1a | 147 ± 7.4 | 41 ± 12.4 | 36 ± 3.5 | 43 ± 1.6 |
| 39 | USP24 | 123 ± 6.3 | 101 ± 13.4 | 78 ± 4.9 | 89 ± 1.7 |
| *40 | FBXL17 | 151 ± 2.1 | 1 ± 1 | 0 ± 0.1 | 0 ± 0.1 |

**Extended Data Table 2 continued: Growth inhibition by individual SIPs represented in Fig. 2b.**
| # | SIP Target | Percent Growth ± SEM | | | |
| --- | --- | --- | --- | --- | --- |
|  |  | Fibro | SK | PHEO | BON |
| *41 | SLBP | 124 ± 4.8 | 2 ± 0.2 | 16 ± 5.1 | 12 ± 1.4 |
| *42 | RIZ | 125 ± 5.3 | 0 ± 0.2 | 0 ± 0.1 | 0 ± 0.1 |
| 43 | RbBP6 | 115 ± 3.4 | 133 ± 10.3 | 88 ± 1.8 | 97 ± 1.7 |
| 44 | USP14 | 114 ± 4.4 | 126 ± 12.4 | 78 ± 6 | 94 ± 1.7 |
| 45 | eIF3eta | 41 ± 2 | 3 ± 1.4 | 9 ± 2.4 | 25 ± 1.2 |
| 46 | Rb1b | 116 ± 3.7 | 57 ± 17.3 | 34 ± 8.9 | 54 ± 3.3 |
| 47 | GPATCH8 | 117 ± 4.8 | 129 ± 9.1 | 79 ± 3.3 | 98 ± 2.2 |
| 48 | ELAVL1 | 110 ± 3.7 | 113 ± 23.3 | 67 ± 7.6 | 70 ± 3.8 |
| 49 | LIN9b | 119 ± 5.4 | 142 ± 11.8 | 78 ± 6.6 | 97 ± 1.9 |
| 50 | ABI1 | 111 ± 3.8 | 91 ± 19 | 52 ± 12 | 82 ± 1.7 |
| *51 | FAM53C | 112 ± 4.7 | 34 ± 11.3 | 57 ± 12 | 61 ± 1.5 |
| 52 | Control 2 (PEN tag) | 112 ± 2.9 | 138 ± 14.3 | 76 ± 5.9 | 94 ± 1.6 |

\* selected as a hit

**Extended Data Table 3:**
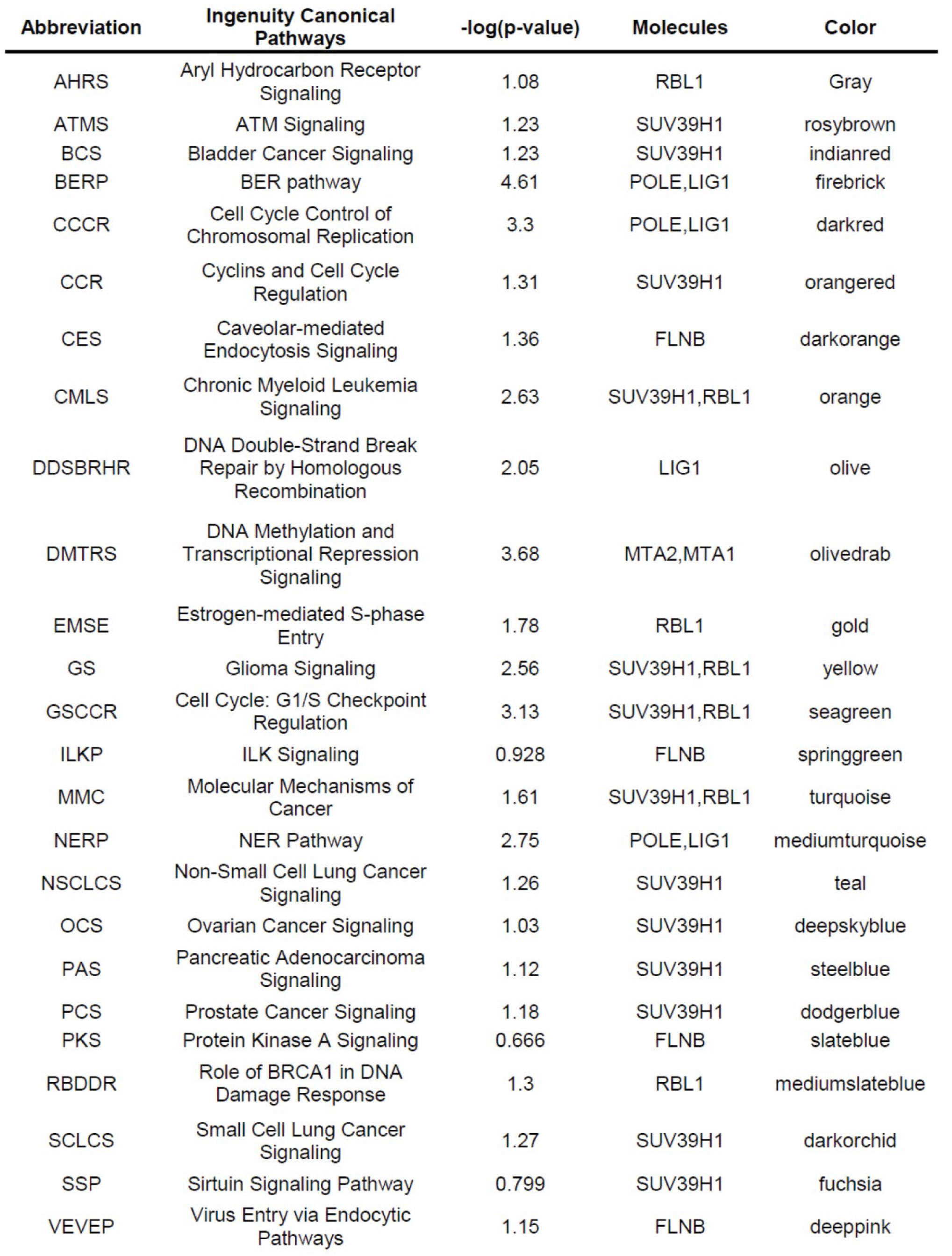
Pathway Enrichment Analysis on 15 selected hits.

**Extended Data Table 4:**
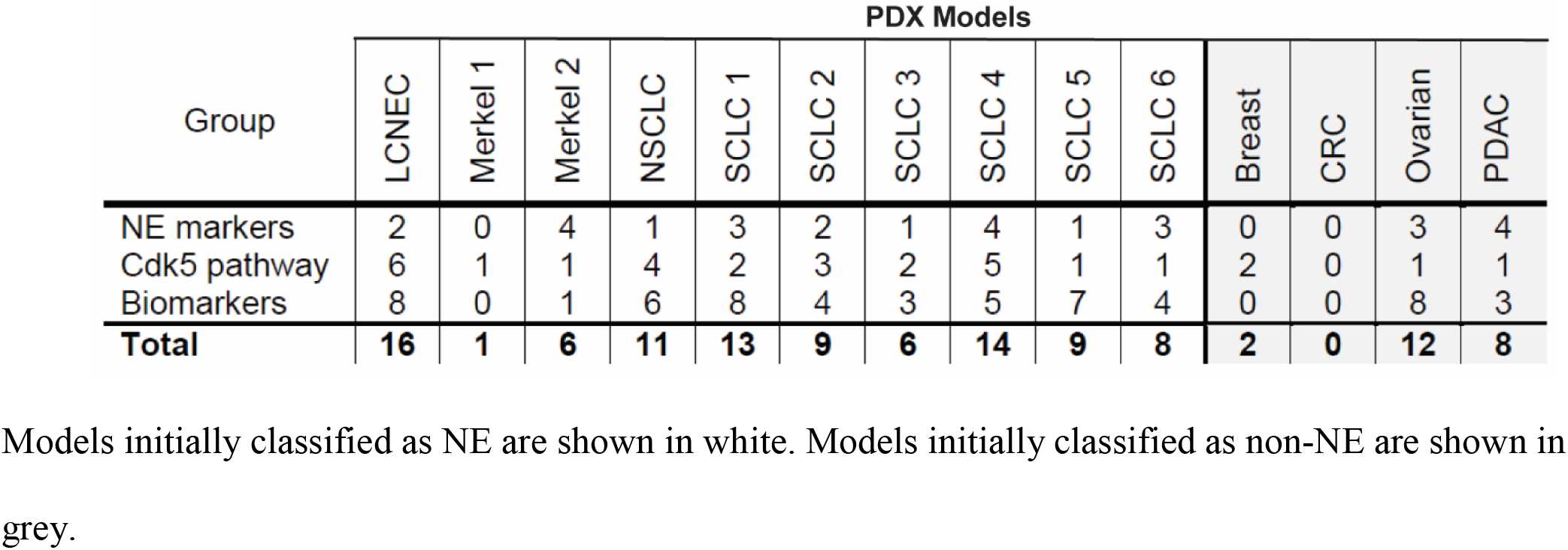
Index Scores for characteristics of PDX models of cancer.

## Notes

https://www.phosphosite.org/Supplemental_Files

